# Evaluation of the antifibrotic potency by knocking down SPARC, CCR2 and SMAD3

**DOI:** 10.1101/376459

**Authors:** Weifeng Ding, Weilin Pu, Shuai Jiang, Yanyun Ma, Qingmei Liu, Wenyu Wu, Haiyan Chu, Hejian Zou, Li Jin, Jiucun Wang, Xiaodong Zhou

**Author notes:** These authors contributed equally to the manuscript (Weifeng Ding, Weilin Pu). Corresponding authors: Xiaodong Zhou, University of Texas-McGovern Medical School, TX, USA., Jiucun Wang, School of Life Sciences, Fudan University, Shanghai, China.

## Abstract

The genes of SPARC, CCR2, and SMAD3 are implicated in orchestrating inflammation and fibrosis in scleroderma and other fibrotic disorders. Aim of the studies was to examine synergistic effect of inhibition of these genes in treating fibrosis. The peptide nanoparticles were used to deliver the siRNAs in bleomycin-induced fibrotic mice. Triple combination of siRNAs targeting on Sparc, Ccr2 and Smad3 achieved favorable anti-inflammatory and anti-fibrotic effects. Inhibition of inflammation was evidenced by reduced inflammatory cells and proinflammatory cytokines in the BALF and/or the tissues. Activation of fibroblasts was suppressed in mouse tissues in which α-Sma and collagens were significantly reduced. Aberrant expression of the genes in fibroblasts, monocytes/macrophage, endothelial and epithelial cells were reinstalled after the treatment. In addition, transcriptome profiles indicated that some bleomycin-induced alterations of multiple biological pathways were recovered to varying degrees by the treatment. The results indicated that the triple combination of siRNAs systemically reinstated multiple biopathways, probably through controlling on different cell types including fibroblasts, monocytes/macrophages, endothelial cells and others. The multi-target-combined therapeutic approach examined herein may represent a novel and effective therapy for fibrosis.

## Introduction

Systemic fibrosis, such as systemic sclerosis (SSc), is often a fetal disease [1]. In US, among the lethal consequence caused by different diseases, the proportion of the fibrosis takes up about 45% [2]. Although various anti-inflammatory, anti-fibrotic and immunosuppressive agents are used to control fibrosis, majority of them have not been proven successful [1, 3, 4]. Therefore, developing novel anti-fibrotic intervention is urgently needed. Pathological prototype of many systemic fibrotic disorders is characterized by excessive extracellular matrix (ECM), inflammation, abnormal immunity and vasculopathy, which are mainly involved in fibroblast, monocytes, lymphocytes and endothelium, as well as the interaction among these different cell types [1, 3].

SPARC (secreted protein, acidic and rich in cysteine) is a matricellular component of ECM mainly expressed by fibroblasts, endothelial cells and lipocytes [5, 6]. It is an important mediator of cell-matrix interaction [7]. It is commonly overexpressed in fibrotic tissue [6, 8–11]. SPARC can stimulate canonical transforming growth factor (TGF)-β pathway [12, 13], and it also plays a possible role in the recruitment of neutrophils to the sites of acute inflammation [14]. Our previous research had demonstrated SPARC inhibition attenuated fibrosis *in vitro* and *in vivo* [15].

CCR2 (C-C chemokine receptor 2) is a common receptor of monocyte chemoattractant proteins (MCP 1,2,3,4 and 5) [16], and it is normally expressed on monocytes, as well as activated T cells, B cells and immature dendritic cells [17, 18]. It not only directs the recruitment of immune cells to the sites of inflammation, but is also involved in angiogenesis, development of fibrosis, migration of fibrocytes to the alveolar space after fibrotic injury [19]. CCR2 and its ligands (MCP1 and MCP3) are highly expressed in SSc-related fibrotic tissues [20]. Blockade of CCR2 ameliorates progressive fibrosis in lung and kidney via suppressing macrophage infiltration and activation [21].

The SMADs (Drosophila mothers against decapentaplegic proteins) are a family of intracellular molecules that act as the main TGF-β signal transducers [22]. As an important member of R-SMADs (receptor-regulatory SMADs), SMAD3 is normally expressed in epithelial cells and lymphocytes [23, 24], and possible other type of cells (e.g. fibroblasts) under disease status or upon TGF-β activation [25]. TGF-β/SMAD signaling plays crucial roles in immune dysregulation, secretion of inflammatory and fibrogenic cytokines and chemokines, and fibroblast activation [26–28]. The Smad3^−/-^ mice are protected against renal tubule-interstitial fibrosis by blocking of endothelial mesenchymal transition (EMT) and abrogation of monocyte influx and collagen accumulation [28].

Overall, SPARC, CCR2, and SMAD3 are three important molecules expressed in various cell types, and they have been implicated in orchestrating the fibrotic response. Though evidences show an individual mediator of fibrosis may have a molecular interplay, a multi-target therapeutic approach may be systemically. Therefore, we selected the siRNAs targeting above three genes and attempted to ameliorate inflammation and fibrosis in mouse models.

## Results

### PNP/PRSsi ameliorated fibrosis in lungs tissues induced by BLM

Low toxicity, high efficiency and low off-target rates of the peptide nanoparticles (PNP, Suppl. Fig. 4) were observed and consistent with the previous report [29]. The expression levels of the targeted genes including *Sparc, Ccr2* and *Smad3* were decreased significantly in the therapeutic groups (Suppl. Fig. 5). In parallel, the expression levels of *Colla1* and *Col3a1* genes of the lung tissues were remarkably reduced in the treatment group with siRNAs for SPARC (P), CCR2 (R), and SMAD3 (S) carried by the PNP (PNP/PRSsi) compared to that in both BLM and Nano (BLM+PNP) groups (Fig. 1A). Moreover, the protein level of type I collagen (Fig 1B) and the soluble collagen content (Fig. 1C) were significantly reduced in the PNP/PRSsi treatment groups compared to those in BLM and Nano groups. The HE (Fig. 1D upper) and Masson Trichrome (Fig. 1D lower) staining showed that the BLM or Nano group had thicker collagen-fiber in the impaired lung alveolar wall compared to those in the saline-injection and the treatment groups. After PNP/PRSsi treatments, only a small quantity of the collagen-fiber was found and the tissue architecture of the lung alveolar was relatively intact (Fig. 1D). These results indicated that the pulmonary fibrosis was significantly alleviated by PNP/PRSsi treatments in the BLM-induced mice.

**Figure 1.**
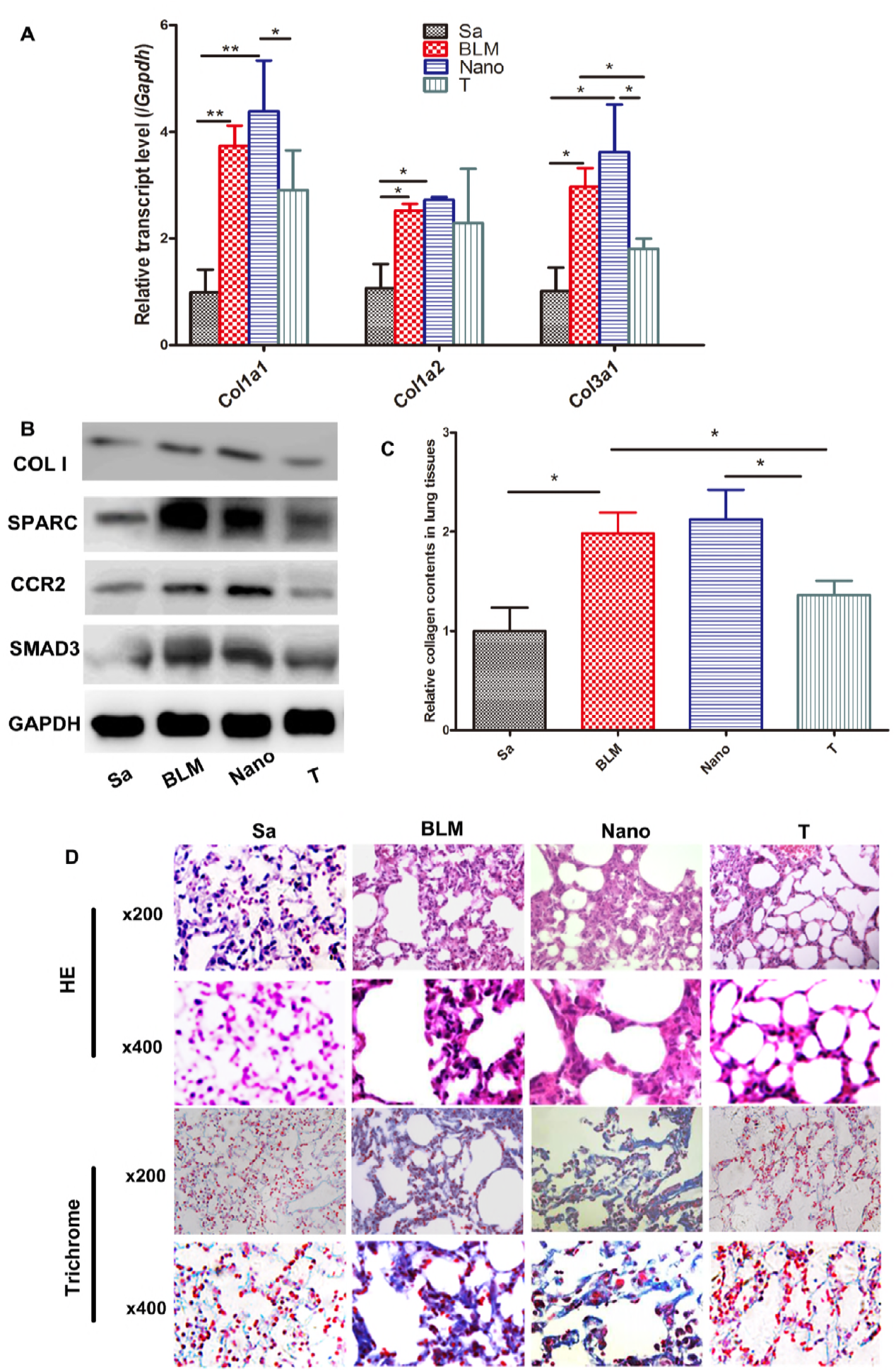
Examination of the anti-fibrotic effect in lungs of mouse models induced by bleomycin (n=4). (**A**) The mRNA levels of collagen *Colla1*, *Colla2*, and *Col3al* were examined by Q-PCR. (**B**) The type I collagen protein was detected by western blotting. COL I: type I collagen protein of the mouse. (**C**) Sircol assays to detect the soluble collagen content in the pulmonary tissues. (**D**) Representative histological figures of HE and Trichrome staining of mouse lung tissues in the different groups. Sa: saline group, injected with saline (negative control); BLM: BLM group, injection with BLM only on day 0; Nano: Nano group, injection with BLM and PNP; T: therapeutic group, injection with PNP/PRSsi; *: *P*<0.05 and **: *P*<0.01.

### PNP/PRSsi antagonized fibrosis in skin tissues induced by BLM

The treatment with PNP/PRSsi significantly reduced the skin *Colla1* and *α-SMA* (alpha smooth muscle actin) mRNA expression (Fig. 2A), as well as type I collagen protein level compared to those in the BLM and Nano groups (Fig. 2B). The soluble collagen content in skin tissues of the mice was also significantly decreased in the therapeutic group (Fig. 2C). On contrast, *Mmp3* (matrix metalloproteinase 3) mRNA expression (Fig. 2D) was remarkably up-regulated in the PNP/PRSsi treatment group. Furthermore, the thickness of the dermis was significantly reduced in the therapeutic groups, in contrast to the BLM or Nano groups that showed increased collagen-fiber bundles and reduced fat tissue in dermal skin by HE and Trichrome staining (Fig. 2E). In addition, the number of detectable capillaries per 100 fold power lens in the dermal skin was significantly increased in the PNP/PRSsi treatment group compared to that in the BLM or Nano groups (Suppl. Fig. 7).

**Figure 2.**
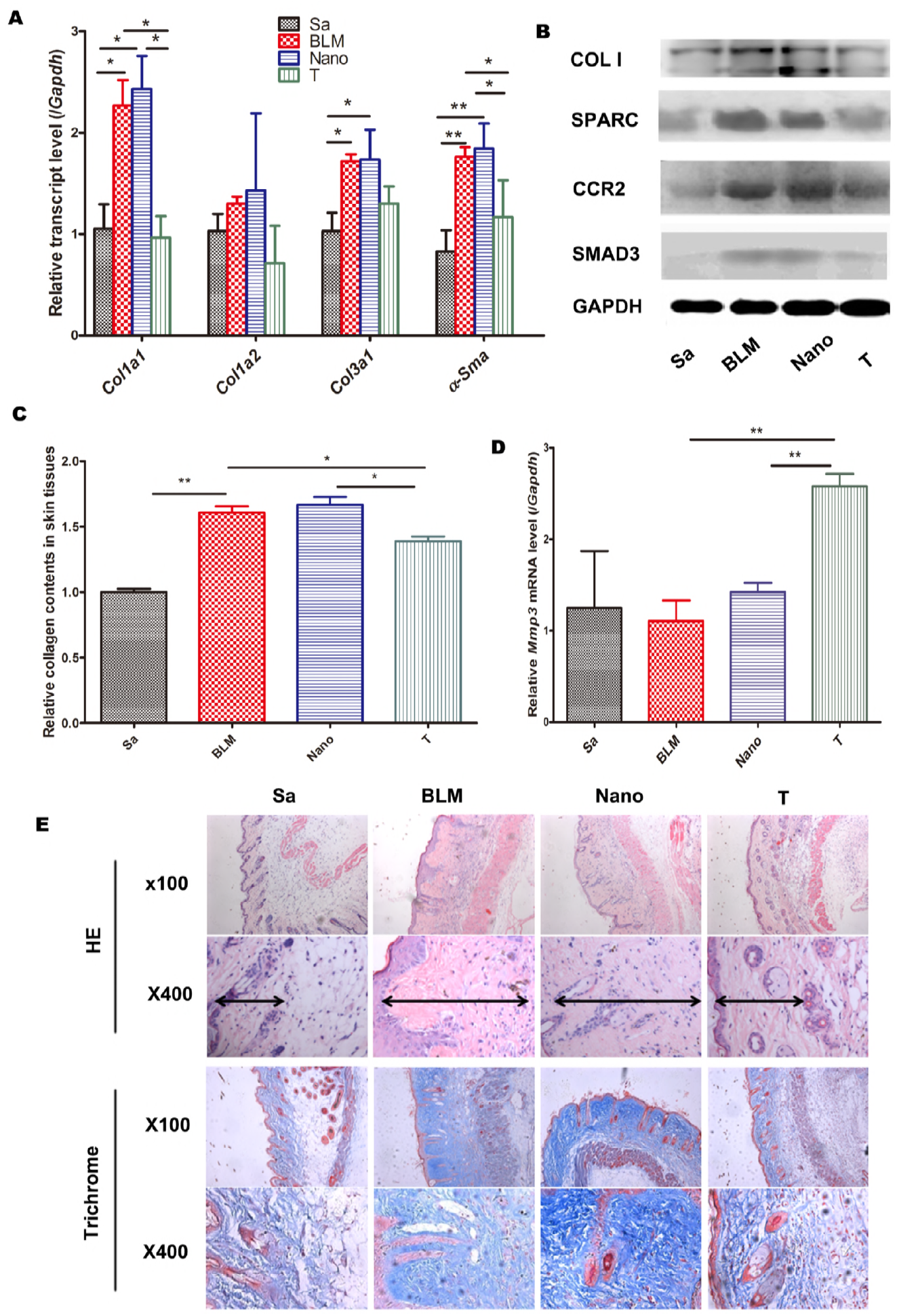
Examination of the anti-fibrotic effect in skin of mouse models induced by bleomycin (n=5) (**A**) The mRNA levels of collagen *Colla1, Colla2, Col3a1*, and *α-SMA* were examined by Q-PCR. (**B**) Collagen I protein was detected by western blotting. (**C**) Sircol assays to detect the soluble collagen content in the skin tissues. (**D**) The mRNA expression level of the anti-fibrotic gene, *Mmp3* was examined by Q-PCR. (**E**) Representative histological figures of HE and Trichrome staining of mouse skin samples in the different groups. Sa: saline group, injected with saline; BLM: BLM group, injection with BLM per day from day 0 to day 21; Nano: Nano group, injection with BLM and PNP per day; T: treatment group, injection with PNP/PRSsi treatments. *: *P*<0.05 and **: *P*<0.01. Di-arrowhead indicated the thickness of the dermis.

### PNP/PRSsi reduced inflammation in the BALF of mouse models with pulmonary fibrosis

To validate the anti-inflammation effect of PNP/PRSsi in the fibrogenesis of lung tissues, we detected inflammatory cytokines and the nucleate cells in the bronchoalveolar lavage fluid (BALF) of the mice. We found that compare to the BLM and Nano groups, the PNP/PRSsi treatment significantly reduced the total number of inflammatory cells, neutrophils and lymphocytes infiltrating as well as actual reduction of macrophages (Fig. 3A and Table 1). Moreover, the concentrations of IL-6 and MMP3 proteins in the BALF were also significantly decreased in the treatment groups (Fig. 3B and 3C).

**Figure 3.**
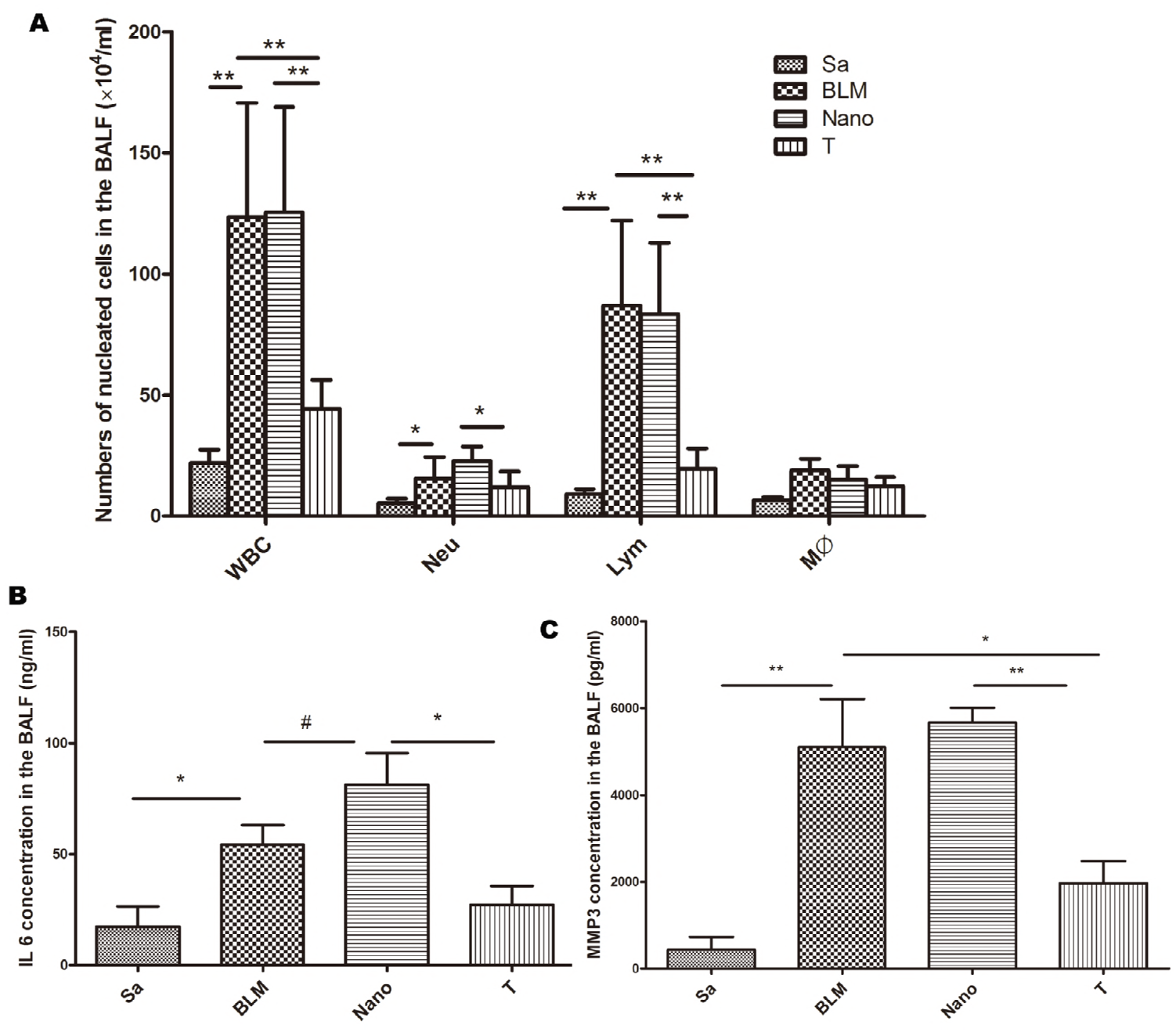
Examination of the inflammatory cells and inflammatory cytokines in the BALF of pulmonary fibrosis mouse models (n=4) (**A**) White blood cells count and differential count by Wright-Giemsa’s staining in the BALF under the optical microscope. The concentration of IL 6 (**B**) and MMP3 (**C**) proteins in the BALF were detected by ELISA. Sa: injected with saline (negative control); BLM: injection with BLM only on day 0; Nano: injection with BLM and PNP; T: therapeutic treatments; WBC, white cells count; Neu, neutrophil; Lym, lymphocyte; Mϕ, macrophage. *: *P*<0.05 and **: *P*<0.01. #, P>0.05.

**Table 1.** The percentage of nucleated cells in the BALF

| Cell<br>count(%) | Sa | BLM | Nano | T |
| --- | --- | --- | --- | --- |
| Neu (%) | 23.2±3.66 | 12.7±7.22 | 20.0±5.73 | 23.7±11.88 |
| Lym (%) | 42.7±3.66 | 68.3±5.51 | 65.6±2.06 | 41.0±11.07* |
| Mac (%) | 31.2±3.36 | 17.0±5.42 | 12.3±1.91 | 34.1±21.81 |

\*: $P < 0.05$ between the T VS BLM or Nano group. Sa: injected with saline (negative control); BLM: injection with BLM only on day 0; Nano: injection with BLM and PNP; T: therapeutic treatments; Neu: neutrophil; Lym: lymphocyte; Mac: macrophage.

### PNP/PRSsi attenuated inflammation and immune response in lung and skin fibrosis mouse models

In the pulmonary fibrotic mouse models, compared with the BLM and Nano groups, the expression level of pro-inflammatory gene *Il-6* was remarkably reduced in the treatment groups (Fig. 4A). *Mmp3* expression was also dramatically decreased when the pulmonary fibrotic mouse models were administrated with PNP/PRSsi (Fig. 4E). The meaning point was that the expression level of *Ifn-γ*, a cytokine with anti-fibrotic activity [30, 31] was up-regulated by application of PNP/PRSsi in the pulmonary fibrotic mouse models, especially in the Nano group (Fig. 4B).

**Figure 4.**
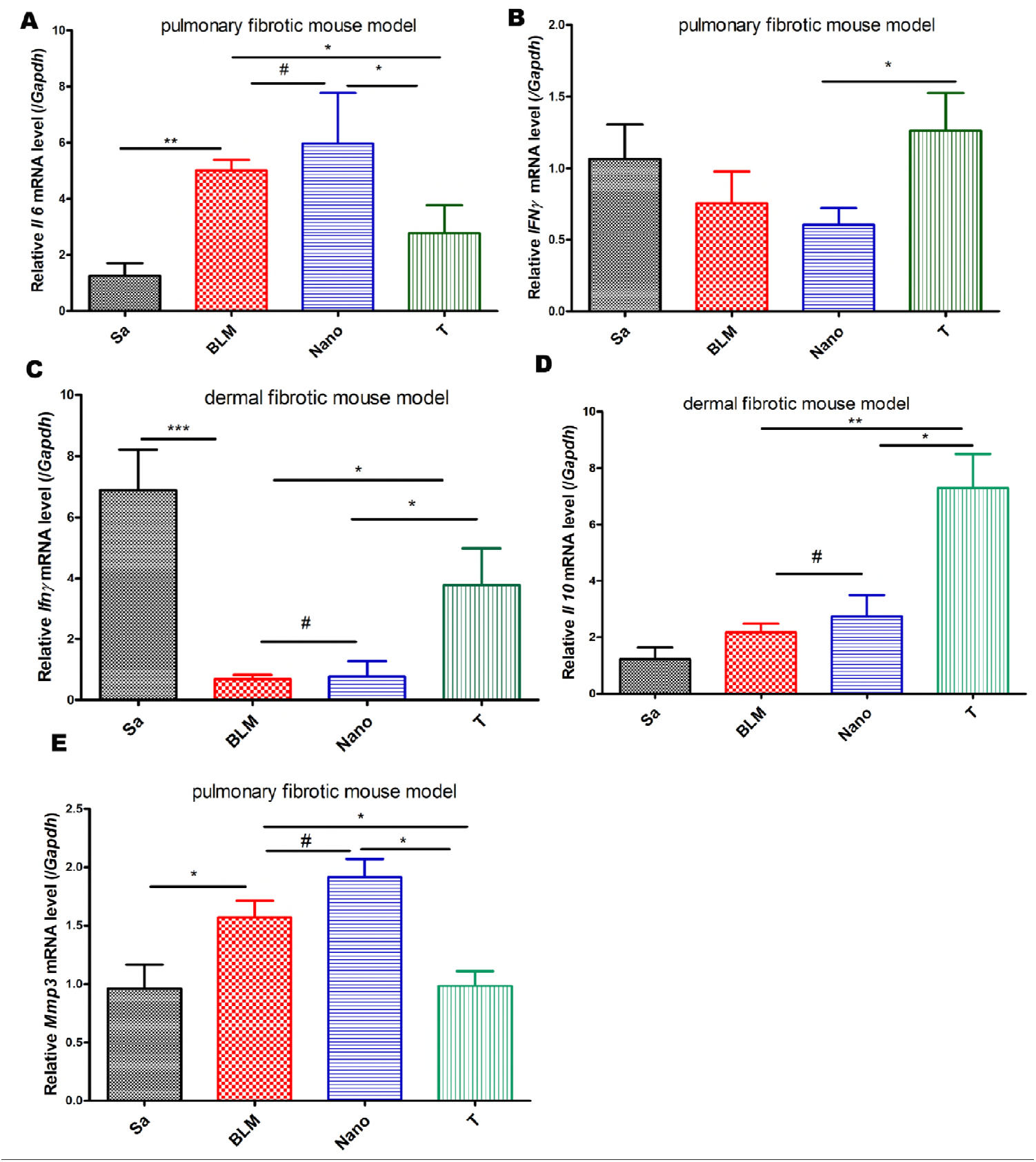
Examination of the cytokines involved in the inflammation and immune response in dermal and pulmonary mouse models by Q-PCR. (**A**) *Il 6* and (E) *Mmp3* mRNA levels were examined in pulmonary fibrosis mouse models. (**B, C**) *Ifn γ* mRNA levels were examined in lung and skin fibrosis models, respectively. (**D**) *Il 10* mRNA levels were examined in dermal fibrosis mouse models. Sa: injected with saline (negative control); BLM: injection with BLM; Nano: injection with BLM and PNP; T: therapeutic treatments. *: *P*<0.05, **: *P*<0.01,***:*P*<0.001. #, P>0.05.

Interestingly, in the mice of skin fibrosis, the observable increase of immunity genes mRNA levels were found to not only include *Ifn-γ* (Fig. 4C) but also involve in the immunity negative regulatory factor *Il-10* (Fig. 4D) after PNP/PRSsi treatment. It implied that PNP/PRSsi could attenuate pro-inflammatory cytokines and also mediated the immunity response via cytokine IL 10 to inhibit fibrosis.

### Gene expression profiles validated the anti-fibrosis effect of PNP/PRSsi in pulmonary model

The heatmap of RNA sequencing based on all of the variable genes were shown in Figure 5A. It showed that the gene expression patterns were similar between saline and PNP/PRSsi treatment groups, while the BLM group had distinct gene expression profile. 156 DEGs (differentially expressed genes) were found between BLM and PNP/PRSsi treatment groups, and 1397 DEGs between BLM and saline treatment groups. Among these DEGs, 103 genes were shared (Fig. 5C). These shared DEGs were mainly involved in Notch signaling (e.g. Notch3, Bcl9l), nuclear receptor signaling (e.g. Rpl39), inflammation (e.g. Mmp9, Ccr5, Mgst1), the SUMO modification (e.g. Sumo1, Ildr2), NOS/hypoxia signaling (e.g.Itga2, Fn1, Ndufa6, Cox6a2), and Toll-like receptor (TLR) pathways (e.g.Tlr9, Oas3) [3, 28, 32–35]. Some of the genes including Notch3, Mmp9, Ildr2, Sumo1, Rpl39, and Ndufa6 also altered in human lung biopsy samples of pulmonary fibrosis with the similar tendency to that in our mouse models, data from GEO database (Suppl. Fig. 8-13). The KEGG enrichment analysis showed most significantly altered biopathways in each comparison of different groups. The top up-regulated ones (e.g. ECM-receptor interaction, focal adhesion, cytokine-cytokine receptor interaction, B cell receptor, protein digestion and absorption) in BLM group were attenuated by PNP/PRSsi treatment group (Fig. 5D). On the contrary, the top down-regulated ones, including metabolic pathways, ribosome, oxidative phosphorylation, cytochrome p450, and drug metabolism in BLM group were partly rescued to the normal status by PNP/PRSsi treatment group (Fig. 5E).

**Figure 5.**
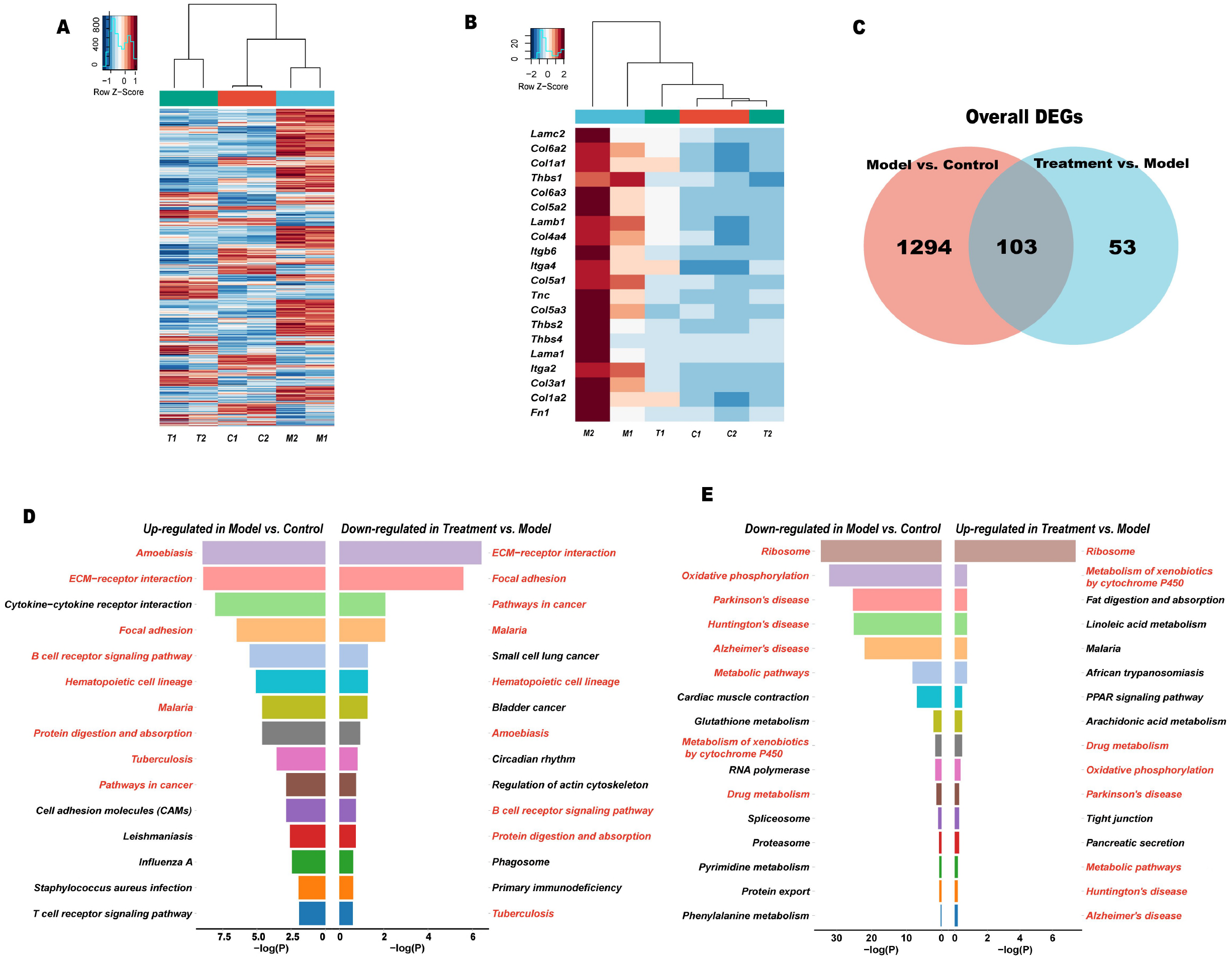
The gene expression profiles in the control, model and PNP/PRSsi treatment mice in pulmonary fibrosis mouse models. (**A**) The heatmap of all the differentially expressed genes (DEGs) among the control, model and treatment groups. Horizontal axis was depicted as a z score of each gene expression level with FPKM (fragments per kilobase of transcript per million fragments mapped). Vertical axis represents each of the DEGs. C: Saline group that the mouse models were injected with saline and served as the negative control, M: the mouse model injected with BLM, which served as a model group, T: the mouse model of treatment group, therapeutic injection with PNP/PRSsi. (**B**) The expression profiles of DEGs in the extracellular matrix (ECM) pathway among three groups were shown. (**C**) The Venn diagram showed the unique and shared numbers of the DEGs in different comparisons. 1294 DEGs were unique in the comparison between model group and control group, while 53 DEGs were unique in the comparison between PNP/PRSsi treatment group and model group. Meanwhile, 103 DEGs were shared between the two comparisons. (**D**) Top 15 significant pathways in KEGG enrichment analysis based on DEGs of different comparisons. Right part represents the down-regulated signaling pathways in the treatment group compared with those of in the model group, while the left part represented the up-regulated signaling pathways in model group compared with those in control group. (**E**) Top 15 down-regulated pathways in BLM group while partly up-regulated in PNP/PRSsi treatment group.

The expression pattern of the genes involved in ECM-receptor interaction pathway was significantly up-regulated in BLM group compared with the saline group, and was then significantly retrieve to normal status after PNP/PRSsi treatment (Fig. 5B). Similarly, the expression profiles of the genes involved in cell adhesion molecules (CAMs) and other pathways (Suppl. Fig. 15) in these three groups showed similar patterns. In addition, we found that the expression levels of genes located in the inflammation-related signaling pathways, including chemokine, NF-κB, and JAK-STAT signaling pathways also showed similar tendency (Suppl. Fig. 16). By contrast, about 25 key genes involved in oxidative phosphorylation and xenobiotics, glutathione metabolic pathways, which were anomaly down-regulated in the model group, were almost revived to the normal status after PNP/PRSsi treatments (Suppl. Fig. 17).

Taken together, the RNA-Seq data indicated that systemic changes of biopathways induced by the BLM were largely recovered by the PNP/PRSsi treatment.

## Discussion

Traditional strategy of anti-fibrosis is to target single gene. However, the target gene is often specific to single biopathway and cell type expression, which may limit its impact to fibrosis that usually involves comprehensive bio-networks conducted by different cell types. In this study, we explored a multi-gene-target approach against fibrosis in BLM-induced mouse model. The three targeted genes are distinct in their primary expressing cells. SPARC is mainly expressed in fibroblast and endothelial cells in which it participates in both TGF-β dependent and independent fibrotic pathways [5, 6, 36], CCR2 in monocyte and lymphocyte involving in inflammatory signaling [17–19], and SMAD3 in lymphocyte and epithelial cells driving canonical TGF-β pathway [23, 24]. These different cell types and pathways are believed to synergistically contribute to systemic fibrotic process, such as in SSc [3, 33, 35, 37, 38]. It is worth noting that like many fibrosis in human diseases, BLM-induced fibrosis in mouse models is TGF-β dependent and inflammation driven process [39–41].

As a result, inhibition of these three target genes attenuated fibrosis in both lung and skin tissues induced by BLM. Underlying molecular changes indicated that the activation of fibroblasts was suppressed as the expression of α-Sma and collagen genes (Fig. 1, Fig. 2, and Suppl. Fig. 14) were generally controlled by the PNP/PRSsi treatment. This improvement appeared in parallel with suppression of inflammation evidenced by reduced inflammatory cells and pro-inflammatory cytokines in the BALF and/or the tissues (Fig. 3 and Fig. 4). While monocytes/macrophage are driven force in inflammation, the CCR2 gene along with other inflammatory genes, such as Chi3l3 and Il1rap, commonly expressed by monocytes/macrophage [42] were significantly restrained after the PNP/PRSsi treatment (Suppl. Fig. 18A). On the other hand, microvasculature in the BLM-induced mouse skin appeared to be improved in the PNP/PRSsi-treated mice as the number of detectable capillaries was significantly increased compared to that in the untreated ones (Suppl. Fig. 7). Some endothelial cell expression genes (e.g. Tiel and Erg) [43] and the gene of Pecam-1 (CD31) that is associated with endothelial-mesenchymal transition (EndoMT) [44] were under controlled in the treated mice (Suppl. Fig. 18B). In addition, some epithelial cell expressing genes (e.g. Clca1 and Serping1) [45] also were recovered from the aberrant expression in the BLM-induced mice (Suppl. Fig. 18C).

A global view of gene expression profile with RNA sequencing analysis indicated that some BLM-induced alterations of major biological signaling pathways, such as ECM, cytokine-cytokine receptor interaction, focal adhesion, metabolic, B cell receptor signaling, protein digestion and absorption pathways were recovered to varying degrees by the PNP/PRSsi treatment. This observation suggested that the PRSsi treatment tended to systemically reconstruct the biopathways dysregulated in the BLM-induced mice.

Among the genes examined, the changes of Mmp3 expression appeared contradictory in skin and lung tissues. It was up-regulated in the former, but down in the latter after the treatment. In fact, MMP3 functions in both anti-fibrotic and pro-inflammatory roles. As an important ECM degrading enzyme, it renders crucial roles in connective tissue remodeling [46, 47]. On the other hand, it may promote the migration and infiltration of the inflammatory cells [48]. Therefore, it is likely that an increased Mmp3 by the treatment may serve as anti-fibrotic function in skin tissue, while a down-regulation could be an indication of controlled inflammation in the lung tissues.

In summary, the application of the combined siRNAs of SPARC, CCR2, and SMAD3 genes ameliorated fibrosis and inflammation in the BLM-induced mice. It systemically reinstated multiple biopathways, probably through controlling on different cell types including fibroblasts, monocytes/macrophages, endothelial cells and others. Considering human fibrotic disorders usually arise from complex networks involving these major cell types, the multi-target-combined therapeutic approach examined herein may represent a novel and effective therapy for fibrosis.

## Materials and methods

### siRNAs

Mouse siRNAs for Sparc and Ccr2 were synthesized by Gene Pharma Inc. (Shanghai, China) and Ribobio Inc. (Guangzhou, China), respectively. Mouse Smad3 Trilencer-27 siRNAs and non-target siRNAs were purchased from Origene Inc. (Rochville, MD, USA). To avoid off-target effect, two pairs of each target gene siRNAs were utilized, and both of them showed the knockdown effect of target genes in a mouse embryo fibroblast NIH 3T3 cell line (Suppl. Fig. 1).

### Animal models of pulmonary fibrosis, delivery of combined siRNAs *in vivo*

Female C57BL/6 mice of about 20 g were purchased from SLRC laboratory animal Inc. (Shanghai, China). Pulmonary fibrosis model was induced by intratracheal instillation of 3 mg/kg bleomycin (BLM, Nippon Kayaku Co., Tokyo, Japan) for one time [41]. A total of 5.6 μg combined siRNAs for Sparc, Ccr2 and Smad3 encapsulated by peptide nanoparticles (PNP) [29] were injected intraperitoneally with 130 μl PBS on day 10, 14, 18 after BLM induction (Suppl. Fig. 2). All groups of mice were sacrificed on day 22 after anesthesia, and the bronchoalveolar lavage fluid (BALF) and the lung samples were collected. The left lungs were ligated for the extraction of BALF, then fixed with 4 % formalin and used for further histological analysis. The right lung tissues were divided into three parts, one for RNA extraction, another one for collagen content analysis, the rest for western blot detection.

### Animal models of dermal fibrosis, delivery of combined siRNAs *in vivo*

For dermal fibrosis model, female C57BL/6 mice about 5 weeks old were induced by local subcutaneous injection of 100 μl BLM (0.5 mg/μl) per day in the shaved lower back for three weeks. The treatments were performed by subcutaneous injection to lower back of 3 μg of combined siRNAs and PNP on day 7, 10, 12, 14, 16, 18, 20 after three hours of BLM administration (Suppl. Fig. 3). All of the mice were sacrificed on day 22 and the skin samples were collected. Five mice were examined in saline group, BLM group, Nano (BLM+PNP) group and therapeutic group, respectively. Saline injections were used as negative controls in both lung and skin fibrosis models. The animal protocols were approved by the Center of Laboratory Animal Medicine and Care of Fudan University.

### Determination of gene expression by quantitative RT-PCR

Total RNA from mouse skin and lung tissues were extracted, complementary DNA (cDNA) was synthesized by using MultiScribe Reverse Transcriptase (Applied Biosystems, Foster city, CA, USA). Quantitative real-time RT-PCR was performed by an ABI 7900 Sequence Detector System (Applied Biosystems). The primers sequences for each gene (*Colla1, Colla2, Col3a1, Sparc, Ccr2, Mmp3, Il-6, Il-5, Il-10, IFN-γ* and *α-Sma*) for SYBR Green I Q-PCR were shown in supplementary table 1 and 2. *Gapdh* gene was served as an internal reference control.

### Detection of protein level by Western blot

Skin and lung tissues were dissolved in RIPA buffer and proteinase inhibitor cocktail (Roche, Basel, Switzerland). The protein concentration was determined by a spectrophotometer using Bradford protein assay kit (Tiangen Biotech Co., Beijing, China). Equal amounts of protein were subjected to SDS-PAGE, then transferred onto 0.45 μm PVDF membranes and incubated with respective anti-mouse primary antibodies, including anti-collagen I antibody (Milipore, Temecula, CA, USA), anti-Smad3, anti-CCR2 antibody (Abcam, Cambridge, MA, USA) and anti-SPARC antibody (Cell Signaling Technology Inc., USA). Anti-GAPDH (Cell Signaling Technology Inc.) was used as an internal control. The secondary antibody was peroxidase-conjugated anti-rabbit mouse IgG. Target proteins were detected with chemiluminescence by using an enhanced chemiluminescence detection system (Thermo, USA). The intensity of the bands was quantified by the Image-QuantTL software (General Electric Company, CT, USA).

### Histological analysis

The tissues were cut into serial 5-μm sections, deparaffinized in xylene and hydrated in a graded series of alcohol. The slides were examined with standard Hematoxylin and eosin (HE) staining to assess the severity of the inflammation and fibrosis of the tissues. The slides were also examined with standard Masson Trichrome staining to evaluate the degree of tissues’ fibrosis.

### Total and differential nucleated cell count in the BALF

BALF samples were taken from the ligated left lung on day 22 after BLM inducing and preserved in buffer containing 1% FBS, 2 U/ml heparin and PBS. Total nucleated cell was counted in duplicate by an optical microscope. Differential cell counts were performed on the thin slides, prepared with smearing BALF samples using Wright-Giemsa’s stain. According to the staining and morphological criteria, differential cell analysis was carried out under a light microscope by counting 100 cells and calculating the percentage of each cell type.

### Determination of concentration of cell factor by ELISA

IL-6 (interleukin-6) and MMP3 concentration from peripheral blood and BALF were determined by using enzyme-linked immuno-sorbent assay (ELISA) according to the manufacture’s protocol (Abcam).

### Determination of soluble collagen content

Non-crosslinked fibrillar collagen in lung and skin samples was measured by the Sircol colorimetric assay (Biocolor, Belfast, UK). Minced tissues were homogenized in 0.5 M acetic acid with about 1:10 ratio of pepsin (Sigma, USA). Tissues were weighted and incubated overnight at 4 °C with vigorously stirring. Digested samples were centrifuged and the supernatant was used for Sircol dyeing to detect the collagen content. The total protein concentration was determined by using Bradford protein assay kit (Abcam) and the collagen content of each sample was normalized to total protein.

### RNA sequencing assay to evaluate the anti-fibrotic effect of multi-genes siRNAs in pulmonary fibrosis mouse models

Total RNA was extracted and used to construct the cDNA library with a KAPA RNA HyperPrepkit (Kapa biosystems, Wilmington, Massachusetts, USA) according to the manufacturer’s instruction. Then the cDNA libraries were sequenced by Illumina HiSeq X Ten systems (Illumina, USA). The average sequencing depth of each fragment was 140 X. The transcriptome data was analyzed with the tophat2 and cufflinks pipeline according to the instructions.

### Statistical analysis

One-way ANOVA (analysis of variance) was used to analyze the difference of multi-groups in treatment. Unpaired t test with welch’s correction was used to test the significance between two groups. A p-value of less than 0.05 was considered as statistical significance.

## Acknowledgements

We declared no competing interests to the manuscript. The work was supported by National Natural Science Founds (grant No. 82318001, 31521003) and Shanghai Municipal Science and Technology Major Project (2017SHZDZX01).

